# Structural Coverage of the Human Interactome

**DOI:** 10.1101/2023.07.24.550328

**Authors:** Kayra Kosoglu, Zeynep Aydin, Nurcan Tuncbag, Attila Gursoy, Ozlem Keskin

## Abstract

Complex biological processes in cells are embedded in the interactome, representing the complete set of protein-protein interactions. Mapping and analyzing the protein structures are essential to fully comprehending these processes’ molecular details. Therefore, knowing the structural coverage of the interactome is essential to show the current limitations. Structural modeling of protein-protein interactions requires accurate protein structures. In this study, we mapped all experimental structures to the reference human proteome. Later, we found the enrichment in structural coverage when complementary methods such as homology modeling and deep learning (AlphaFold) are included. We then collected the interactions from the literature and databases to form the reference human interactome resulting in 117,897 non-redundant interactions. When we analyzed the structural coverage of the interactome, we found that the number of experimentally determined protein complex structures is scarce, corresponding to 3.95% of all binary interactions. We also analyzed known and modeled structures to potentially construct the structural interactome with a docking method. Our analysis showed that 12.97% of the interactions from HuRI, 73.62%, and 32.94% from the filtered versions of STRING and HIPPIE could potentially be modeled with a high structural coverage or accuracy, respectively. Overall, this paper provides an overview of the current state of structural coverage of the human proteome and interactome.

**Significance Statement:** We gathered binary protein-protein interactions from three prominent interactome databases to create a comprehensive human reference interactome. We quantified the structural coverage of the human interactome using already available structural data from four different sources. We further evaluate the percentage of interactions that can be accurately predicted using docking methods.

## Introduction

Protein interactions (PPIs) are key players in many cellular processes [1, 2]. Constructing a complete and accurate interactome (i.e., the network of protein-protein interactions) is crucial to understand better fundamental working principles of cells, functions, and disease mechanisms and eventually to identify key proteins or pathways for developing new treatment strategies. Several experimental and computational studies aimed to determine interactomes and were released either as resources or software [3–19]. These resources are curated and integrated to eliminate experimental artifacts and false positive interactions, yielding up to millions of PPIs. Still, the number of studies incorporating the ever-increasing 3-dimensional (3D) protein structures to the interactome has been limited, posing an ongoing challenge. Structurally characterized interactomes are essential to find detailed proteome level functional annotations [20]. They also help infer the dynamics of the interactions, such as distinguishing permanent versus transient interactions, simultaneously occurring and mutually exclusive interactions, and the temporal order of the interactions [21]. Presence of mutations in protein-protein interfaces [22] and their impact as gain- or loss-of-function can also be revealed with structural information [23]. Drug design and repurposing similarly require structural characterization [24]. Overall, structural annotation of the PPI networks is necessary for molecular-level comprehension of the human interactome.

Understanding the atomic-level interactions between two proteins and the specific amino acids involved is essential for comprehending PPIs at the molecular level. In our previous studies, we developed the PRISM algorithm that uses known protein interfaces to accurately predict structural complexes of protein interactions [25, 26] and a prediction model for hotspots at protein interfaces, which are potential drug targets [27–29]. Other computational methods that are present, and not limited to, are Interactome3D [4] and Interactome INSIDER [3]. Predictions from these studies were further used for elaborating the impact of mutations [30], structural modeling of signaling pathways [31–35], and analysis of post-translational modifications. The predictive performance of these computational methods highly depends on the completeness of the proteome and the availability of protein structures. Human proteome is now 93.2% complete [36], and structural data is dramatically boosted with the accumulated experimental data (Protein Data Bank (PDB) [37]), homology modeling (ModBase [38], SWISS-MODEL [39]) and models from deep-learning methods such as AlphaFold [40]. AlphaFold is also adapted for modeling protein complex structures (i.e., AF2Complex) [41]. Other methods, such as AF2 followed by FoldDock, report promising results. However, these methods may have varying prediction performances on different organisms, and be affected by the size of the protein complexes and post-translational modifications [42, 43]. Despite these advances, AlphaFold’s contribution to modeling the human structural interactome is still relatively limited, estimated to be less than 5% [44].

In this study, we assess the structural coverage of human proteome and interactome by considering available known and predicted protein structures. We first estimated the experimental structural coverage of the reference human proteome. Next, we showed the improvement of structural proteome coverage when complementary methods like homology modeling (SWISS-MODEL, ModBase) and deep learning (AlphaFold) were utilized (Figure 1a). We further assessed the structural coverage of the reconstructed interactomes (obtained by combining STRING, HuRI, and HIPPIE and filtered to produce a comprehensive list of protein-protein interactions). Proteome analysis followed by interactome analysis allowed us to identify the portion of the human interactome that can be predicted using structure-based techniques. This assessment involves determining the proportion of the interactome for which there are complete structural models for both interactors (Figure 1b).

**Figure 1.**
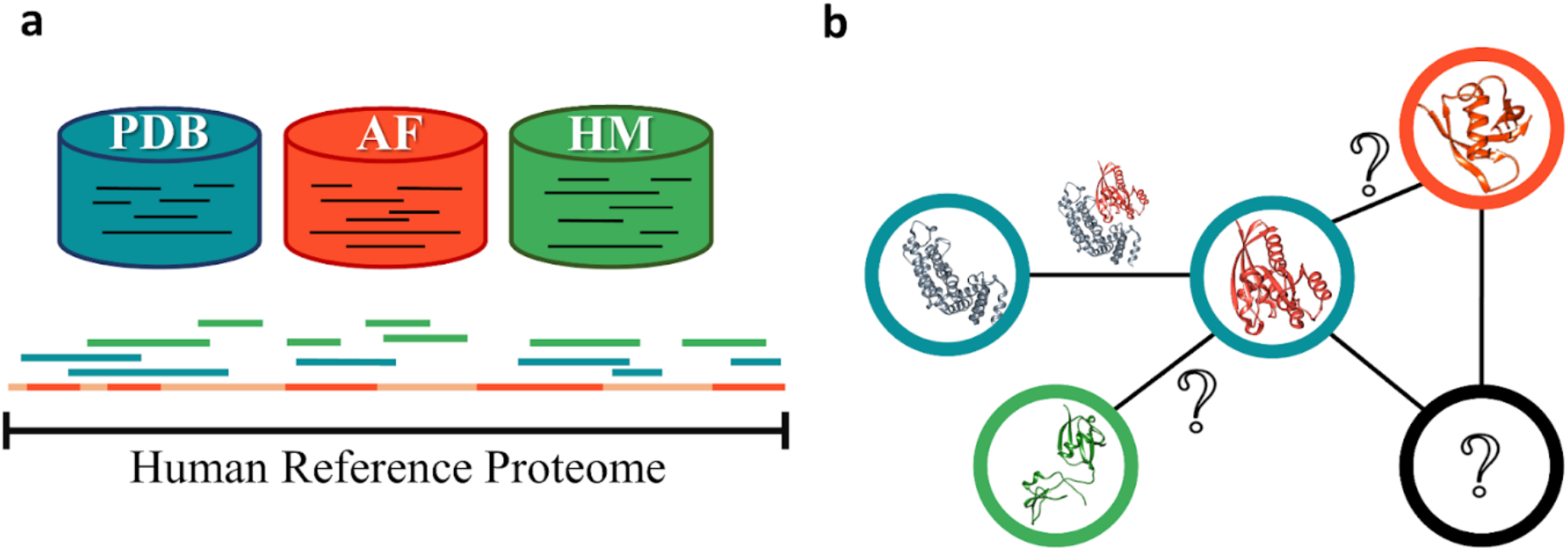
Concept figure. Each database is represented with the same coloring code in both sections. Blue represents sequences from PDB, orange AlphaFold and green homology modeling databases (ModBase and SWISS-MODEL) (a) for reference human proteome. HM: homology modeling, AF: AlphaFold. Human reference proteome is shown by a long black line. Homology models and PDB structures might have overlapping regions that are represented by discrete colored lines. While AlphaFold provides a model for all of the proteome (light red), we focus on regions that are modeled with high accuracy (dark red). (b) Sample network representation of the reference interactome. Protein structures represented by nodes and interactions by edges. Question marks within the nodes are showing monomers that do not have any known 3-D structure in any of the databases. Question marks on the edges show unknown 3-D structures of the interactions (complex) between structurally known monomers.

This work assesses all existing 3-D structural data mapped to human interactome with stringent filtering. These statistics reflect how close we are to constructing the complete structural interactome through experimental and computational methods.

## Results

### Predictive methods improve structural coverage of human proteome

The human reference proteome has 18,401 proteins, of which 7,085 are fully or partially covered in PDB (see methods for coverage calculations). Our results are based on PDB structures containing at least 30 consecutive residues in the corresponding proteins with missing coordinates discarded for each PDB file, and sequence identity with a PDB chain is 100%. At residue resolution, the human reference proteome has 10,789,741 residues, of which 2,125,738 residues have coordinates in PDB that correspond to 19.70% of the proteome (**Table 1**) consistent with previous studies [45]. Sequence coverage categories at different coverage intervals and corresponding percentages of available structures in human reference proteome are given in **Table 2**. Calculations showed that 1,663 proteins, which correspond to ∼9.93% of the reference proteome, are at least 90% structurally covered by PDB. This result indicates that only a small subset of the experimental structures is available for applications that require detailed information, such as molecular simulations, structure-based drug design, and structural interactome construction. Additional computational methods, including homology-based and AI-based structural modeling, are required to expand the set of structurally known proteins.

**Table 1.** Residue-based coverage percentages of whole reference proteome by structure databases.

|  | **PDB** | **SWISS-MODEL**<br>**(NA in PDB)** | **SWISS-MODEL** | **ModBase**<br>**(NA in PDB)** | **ModBase** |
| --- | --- | --- | --- | --- | --- |
| % coverage<sup>#</sup> | 19.70 | 10.09 | 24.99 | 11.95 | 22.18 |

<sup>#</sup> The number of amino acids where structural data is available is divided by the total number of residues in the reference proteome. Overlapping regions are counted only once, and missing residues are not included in the calculation.
NA in PDB: not available in PDB

**Table 2.** Protein-based coverage/accuracy percentages_#_ of reviewed proteins in reference proteome by structure databases.

| **Protein Residue**<br>**Coverage**<br>**(%)** | **PDB**<br>**(%)** | **SWISS-MODEL**<br>**(%)** | **ModBase**<br>**(%)** | **AlphaFold \* (%)** |
| --- | --- | --- | --- | --- |
| ≥ 90 | 9.93 | 13.59 | 17.12 | 17.04 |
| ≥ 70 | 19.25 | 25.99 | 26.07 | 52.52 |
| ≥ 50 | 24.85 | 32.53 | 31.10 | 74.66 |
| < 50 | 13.66 | 10.33 | 15.73 | 23.61 |

<sup>#</sup> For each protein, available structural data are combined. Coverage percentage is calculated by dividing the number of proteins that are modeled above an arbitrary threshold to the total number of proteins in the reference proteome which is 18,401. Overlapping regions are counted only once, and missing residues are not included in the calculation.
\* For AlphaFold, accuracy was calculated instead of coverage.

For the proteins in the human reference proteome, we obtained high-quality models with 30 or more residues from the SWISS-MODEL and ModBase databases, resulting in 7,886 proteins via SWISS-MODEL and 8,618 proteins via ModBase. A total of 11,140 unique proteins were modeled, with 5,364 proteins having predicted models in both SWISS-MODEL and ModBase.

SWISS-MODEL and ModBase provided 24.99% and 22.18% residue-based coverage of the human proteome, respectively, both higher than the reported 19.70% coverage of PDB. The residue-based proteome coverage of the combination of these databases corresponds to 33.02% of the human reference proteome. These results show that homology modeling can cover approximately one-third of the human proteome without any contribution from PDB.

To assess the contribution of homology models to reference proteome coverage, we excluded residues already covered by existing PDB structures and only used residues covered by homology modeling databases. We found 4,904 proteins for SWISS-MODEL and 7,300 proteins for ModBase. A total of 9,096 unique proteins were modeled, with 3,108 proteins having predicted models in both SWISS-MODEL and ModBase.

Residue-based coverage of the proteome unavailable in PDB is 10.09% by SWISS-MODEL and 11.95% by ModBase. Residue-based coverage of their combination shows that homology models contribute to PDB structures by increasing human reference proteome coverage by 16.47%.

Next, we utilized the AlphaFold database produced by an AI-based method (AlphaFold v2.0). The accuracy of the predicted models is provided for each residue with a pLDDT score representing the per residue estimate of its confidence. Residue positions with pLDDT ≥ 70% were considered high-quality [40]. Residue-based proteome coverage by AlphaFold showed that 58.26% of residues have pLDDT ≥ 70%. For structural coverage, we labeled a protein as “accurately predicted” if 85% of its residues were predicted with ≥ 70% pLDDT score. This constraint resulted in the loss of ∼75% of the predicted structures in reference human proteome. Only 4,930 (26.79%) of the AlphaFold predictions satisfy this condition. Among these, 2,425 proteins already have at least a PDB structure, and the remaining 2,505 predictions do not have any known structures deposited in any database before. As previously stated, 7,085 proteins are fully or partially covered in PDB. All PDB data and accurately predicted AlphaFold models combined represent 9,590 proteins (52.11%) for the human proteome.

We further assessed if AlphaFold’s prediction accuracy varies depending on the predicted protein class. We selected five protein classes: enzymes, antibodies, membrane proteins, transcription factors and transporter proteins. We observed that transcription factors have the lowest count in terms of having a PDB structure and the lowest coverage. Likewise, AlphaFold predicts almost all the proteins, 1,451 out of 1,459, within the transcription factor class, yet only 19 passed our high accuracy thresholds. Similarly, AlphaFold predicts all 74 proteins belonging to antibodies. However, it only predicts 17 of them with high accuracy, showing that the prediction accuracy is positively correlated with structures deposited in the PDB and significantly varies between the classes (**Table 3**). In summary, despite the high number of predicted proteins by AlphaFold, the high accuracy predictions are one-quarter of the total protein counts within the class for almost all protein classes.

**Table 3.** Structural data according to protein classes.

| **Functional class** | **Total protein count** | **Proteins [Have PDB]**<br>**(Average PDB coverage, %)** | **Protein count [All predictions w/ AlphaFold]**<br>**(Average pLDDT score, 1-100)** | **Protein count [High acc. predictions w/ AlphaFold]**<br>**(Average pLDDT score, 1-100)** |
| --- | --- | --- | --- | --- |
| Enzymes | 3,618 | 2,114 (72.2) | 3,574 (83.5) | 1,831 (91.6) |
| Antibodies | 74 | 25 (85.6) | 74 (89.6) | 17 (90.5) |
| Membrane Proteins | 5,013 | 1,500 (60.5) | 4,939 (78.1) | 1,391(88.8) |
| Transcription Factors | 1,459 | 372 (30.7) | 1,451 (63.2) | 19 (86.9) |
| Transporter Proteins | 1,843 | 721 (67.8) | 1,823 (78.6) | 517 (88.3) |

Our results prove that combining PDB structures with homology and AlphaFold models increases the structural coverage (**Table 2**). In **Figure 2a**, we show the number of proteins in the human reference proteome shared by the structure databases when all accurate structures are considered. ModBase contributes the most by providing structures exclusively for 1,599 proteins. Also, in **Figure 2b**, we show how the partial and complete coverage changes. PDB covers 1,828 proteins (9.93%) when only highly covered (90%) structures are considered. The addition of high coverage (≥ 90%) homology models and accurately predicted (85% of its residues were predicted with ≥ 70% pLDDT score) AlphaFold models raise this percentage to 33.16%.

**Figure 2.**
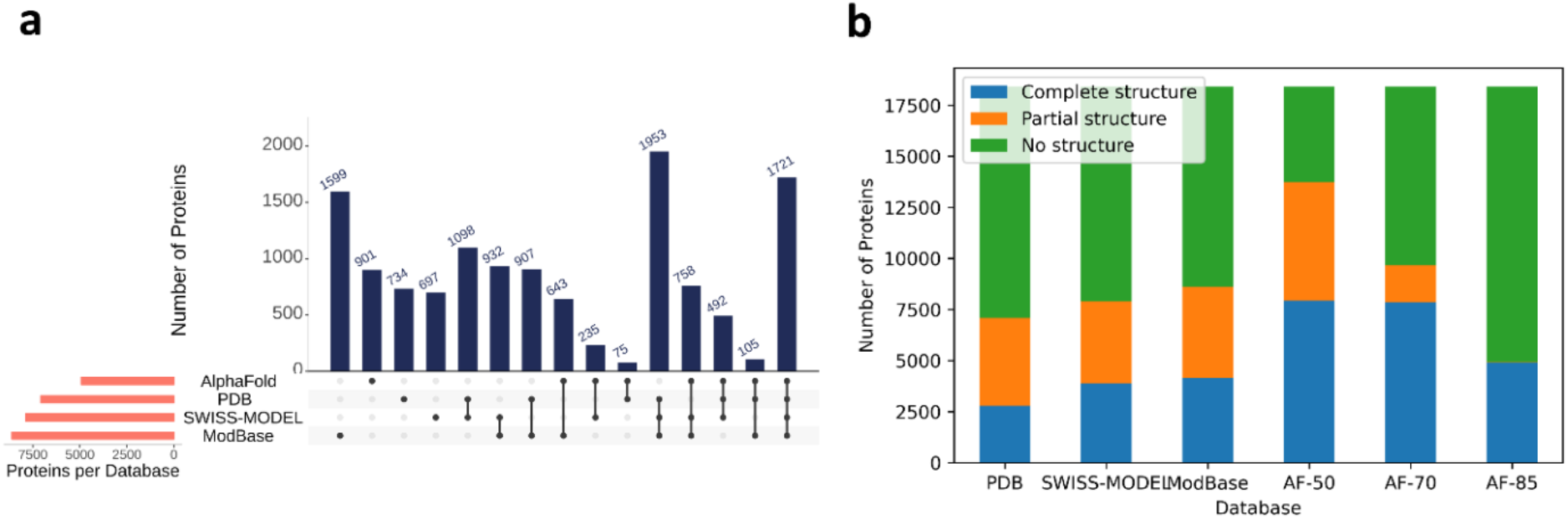
Structural coverage of proteins in the human reference proteome by databases. (a) The number of proteins modeled by PDB, SWISS-MODEL, ModBase, AlphaFold and their intersections are visualized. AlphaFold models with 85% of their residues predicted with ≥ 70% pLDDT score are used. (b) The number of proteins with no structure, partial structure, and complete structure. AlphaFold models with 85% (AF-85), 70% (AF-70), and 50% (AF-50) of their residues predicted with ≥ 70% pLDDT score are used for this demonstration. Partial structure denotes structure coverage < 80% for PDB and homology models. For AlphaFold, it denotes an average accuracy of < 80%. Similarly, complete structure means ≥ 80% coverage or average accuracy. Although it’s not visible, AF-85 has 13 models with partial structures.

### A reference human interactome can be constructed by integrating multiple resources

Estimating the exact size of the human interactome still remains challenging [15]. Available interactions in databases, obtained via multiple techniques, are the best resource for reconstructing a complete human interactome. Here, we analyzed eight major human interaction databases, HuRI, STRING, BioPlex, BioGRID, HIPPIE, IID, APID, and PICKLE, to evaluate the current status of the interactome coverage with available structural data and decided on which database to use toward the construction of a comprehensive structural human reference interactome. We investigated the number of proteins and interactions available after mapping the interactions to the human reference proteome and removing redundant interactions with the same UniProt identifier. Then, we quantified the number of proteins and interactions in each database with a structure and/or a model available in our structural data sources. These statistics are summarized in **Table 4**. HuRI is one of the most comprehensive experimental data providing direct physical interactions for 48,763 PPIs. The filtered STRING database, hereinafter referred to as STRING_F_, resulted in 57,192 physical interactions with high confidence scores (>0.7), considering only experimental and database channels. In addition, there are 53,136 interactions in the BioPlex database. However, these are not restricted to binary physical interactions because the affinity purification–mass spectrometry (AP-MS) technique finds all physical/non-physical interactions in a complex [46]. Lastly, filtering the HIPPIE dataset, after referred to as HIPPIE_F_, to binary high-confidence interactions resulted in 22,280 PPIs.

**Table 4.** Statistics of the number of proteins and protein-protein interactions in interactome databases that are mapped to human reference proteome.

| Database | total number of |  | number of proteins with available structures |  |  |  | number of interactions with structures available for both interacting proteins |  |  |  |
| --- | --- | --- | --- | --- | --- | --- | --- | --- | --- | --- |
|  | interactions | proteins | PDB | SWISS-MODEL | Mod Base | AF* | PDB | SWISS-MODEL | Mod Base | AF* |
| HuRI | 48,763 | 7,889 | 3,317 | 3,414 | 3,729 | 2,026 | 7,046 | 7,014 | 8,938 | 2,737 |
| BioPlex v3.0 | 53,136 | 8,806 | 3,888 | 4,242 | 4,615 | 2,819 | 14,036 | 13,945 | 15,162 | 5,802 |
| STRING v11.5 | 57,192 | 7,327 | 4,518 | 4,214 | 4,095 | 2,085 | 38,005 | 24,829 | 21,442 | 9,941 |
| HIPPIE v2.3 | 22,280 | 7,640 | 4,164 | 3,971 | 4,231 | 1,910 | 10,193 | 7,795 | 8,412 | 1,586 |
| APID v2021_03 | 125,722 | 14,854 | 6,508 | 6,774 | 7,236 | 3,897 | 39,922 | 32,453 | 36,390 | 8,394 |
| PICKLE v3.3 | 211,943 | 15,922 | 6,852 | 4,349 | 7,719 | 4,191 | 88,408 | 68,328 | 71,761 | 14,794 |
| BioGRID v4.4.216 | 719,566 | 17,100 | 6,914 | 7,499 | 8,174 | 4,587 | 292,081 | 233,361 | 239,226 | 70,455 |
| IID v2021_05 | 542,157 | 17,331 | 7,015 | 7,636 | 8,250 | 4,600 | 235,632 | 181,337 | 185,030 | 53,251 |

### Assessment of human interactomes from HuRI, HIPPIE_F_ and STRING_F_: Mapping experimental and predicted 3D structures

**Figure 3** shows the total number of interactions and the interactions with structures/models listed in **Table 4**. We chose two databases, HuRI and BioPlex, which output the results of major experimental studies, as well as the well-known databases STRING which enables filtering the physical interactions by a confidence score, and the HIPPIE which can be filtered with respect to experiment type and confidence score. We demonstrate that there is little overlap in terms of interactions among these four major databases (**Figure 3a**). **Table 4** shows that the HuRI, HIPPIE_F_, and STRING_F_ interaction networks contain 7889, 7640, and 7327 proteins, respectively. While most of the proteins are present in all three databases, a considerable number of proteins are exclusively present in one (**Figure 4**). We also found the number of interactions where both partners had PDB structures. The results show that STRING_F_ has the highest PDB coverage among interactome databases, while HuRI and HIPPIE_F_ have poor PDB coverage (**Figure 3b**). The striking disparity must be due to ribosomal and mitochondrial proteins being significantly less represented in HuRI (protein count: 125; interaction count: 1,348) and HIPPIE_F_ (protein count: 186; interaction count: 2,175) compared to STRING_F_ (protein count: 1,801; interaction count: 16,453), taking part in so many interactions. Given that the Y2H technique detects the bulk of HIPPIE_F_ interactions and all HuRI data, the missing proteins may be due to Y2H’s technical limitations.

**Figure 3.**
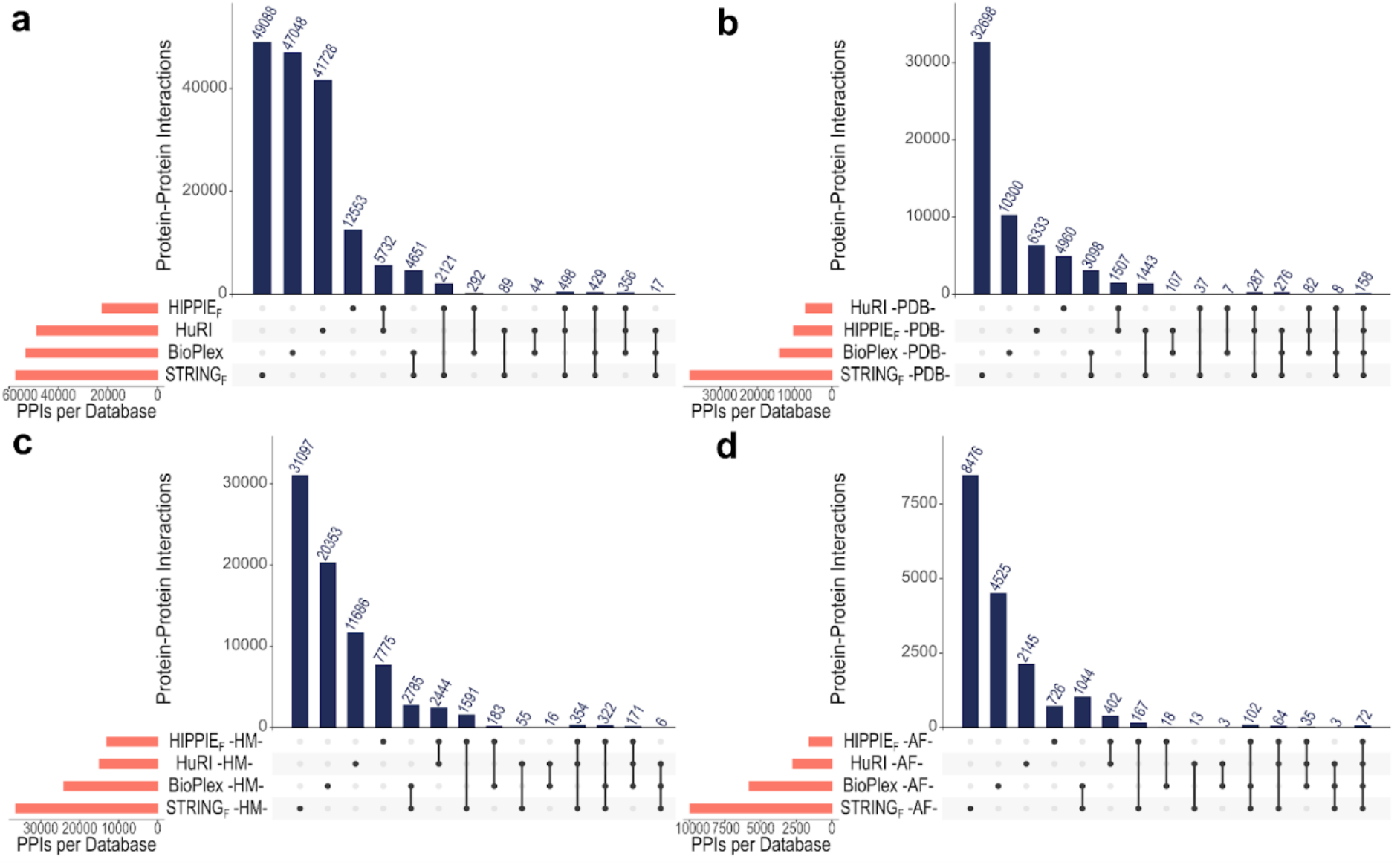
Overview of PPIs found in HuRI, BioPlex, HIPPIE_F_, and STRING_F_ databases. (a) Interactions are filtered according to the reference proteome. (b) Interactions with a PDB structure for both interacting proteins filtered to the reference proteome. (c) Interactions with a high-quality homology model for both interacting proteins filtered to the reference proteome. (d) Interactions with an AlphaFold model that have 85% of their residues predicted with ≥ 70% pLDDT score for both interacting proteins filtered to the reference proteome. HM: homology modeling, AF: AlphaFold.

**Figure 4.**
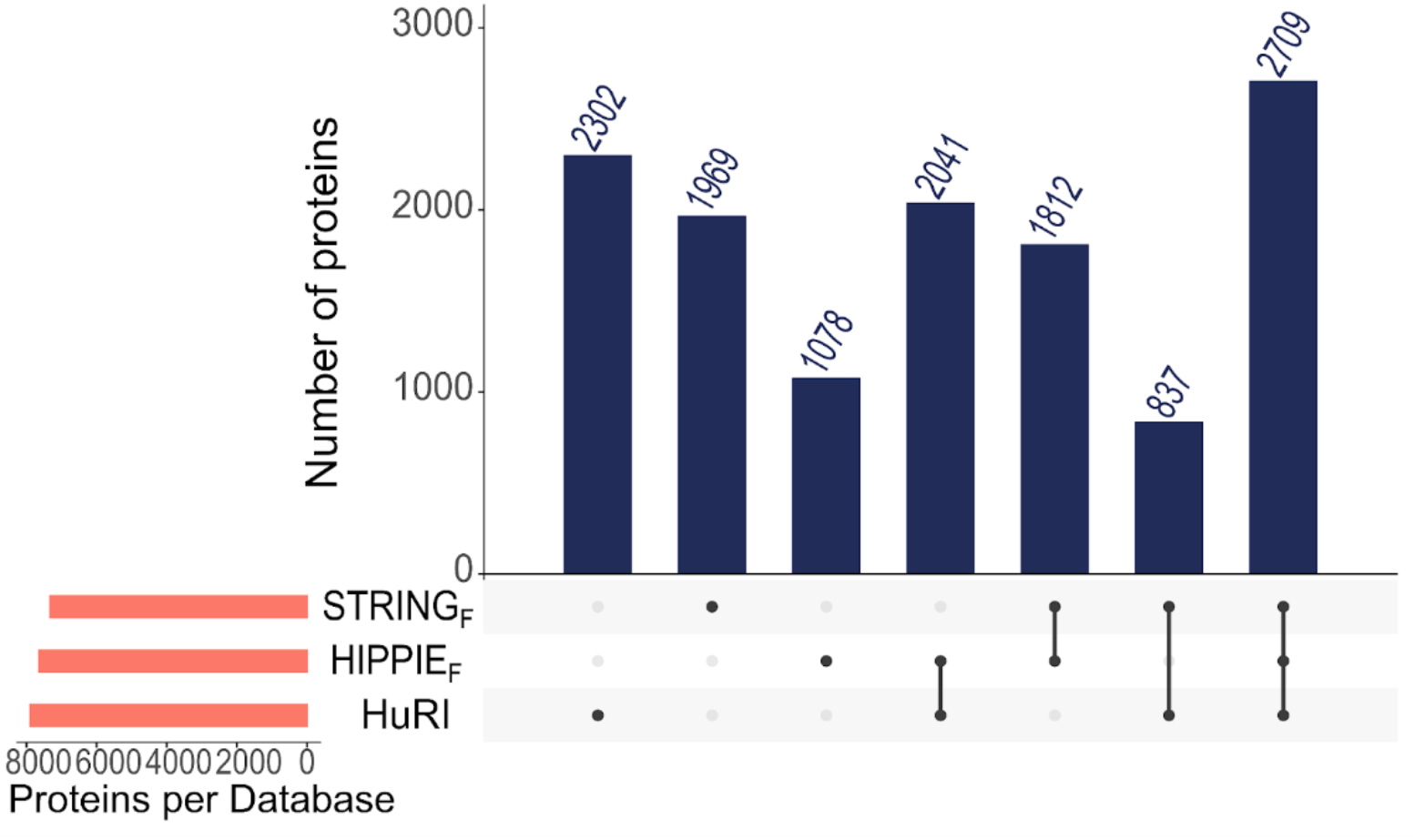
Total number of proteins found in STRINGF, HIPPIEF and HuRI databases.

**Figures 3c and 3d** show the number of interactions with homology and AlphaFold models for both interacting proteins, respectively. Homology models have better interaction coverage compared to AlphaFold. STRING_F_ has the most coverage in both scenarios, while HIPPIE_F_ has the least. In general, the overlap between databases is minimal, reinforcing the notion that integrating these databases is critical since relying on a single database may result in incomplete interactomes.

Since many human proteins are multi-domain, and interactions may occur between single-domain and multi-domain proteins, we analyzed the differences across various interactome databases. The domain count distribution of interacting proteins, as depicted in **Table 5**, reveals an interesting trend. Most interactions occur between single-domain proteins, as evidenced by the significantly high number of interactions in the HuRI interactome, where 19,210 out of 41,705 interactions fall into this category. This observation suggests that single-domain–single-domain protein pairs predominantly mediate interactions in the HuRI dataset may be attributed to potential limitations in correctly expressing lengthy multi-domain proteins in yeast [47], which could result in an underrepresentation of interactions involving these proteins in the dataset.

**Table 5.** Classification and observance percentages of domain types of the interacting protein pairs found in reference interactomes.

|  |  | Single—Single | Multi—Multi | Single—Multi |
| --- | --- | --- | --- | --- |
| Reference Interactomes | HuRI (41,705) | 19,210 | 5,403 | 17,092 |
|  | STRING (54,631) | 19,754 | 12,541 | 22,336 |
|  | HIPPIE (20,601) | 5,445 | 6,248 | 8,908 |

These results have led us to select the HuRI, STRING_F_, and HIPPIE_F_ databases as our reference interactome sources. When these three interactomes are combined, 12,748 non-redundant proteins participate in 117,897 interactions. Mapping these interactions to experimental 596,919 binary protein-protein complexes (https://github.com/ku-cosbi/ interactome-structural-coverage/blob/main/data/PDB_interface_data.tsv) from PDB (as of December 2022) showed that very few experimentally resolved protein interactions are available and they account for only 3.95% of all non-redundant binary interactions. Next, we concentrated on the interactions and displayed the structural coverage of protein-protein interactions in interactome databases (**Figure 5**). As can be seen from the percentages, the STRING_F_ database outnumbers the rest (blue, orange, and green in the inset plot). Experimental and modeled structures of both interactors in HuRI are available for only ∼40% of the whole interactome. In comparison, this figure rises to ∼85% in STRING_F_. Here, we did not distinguish between high or low coverage/accuracy structures.

**Figure 5.**
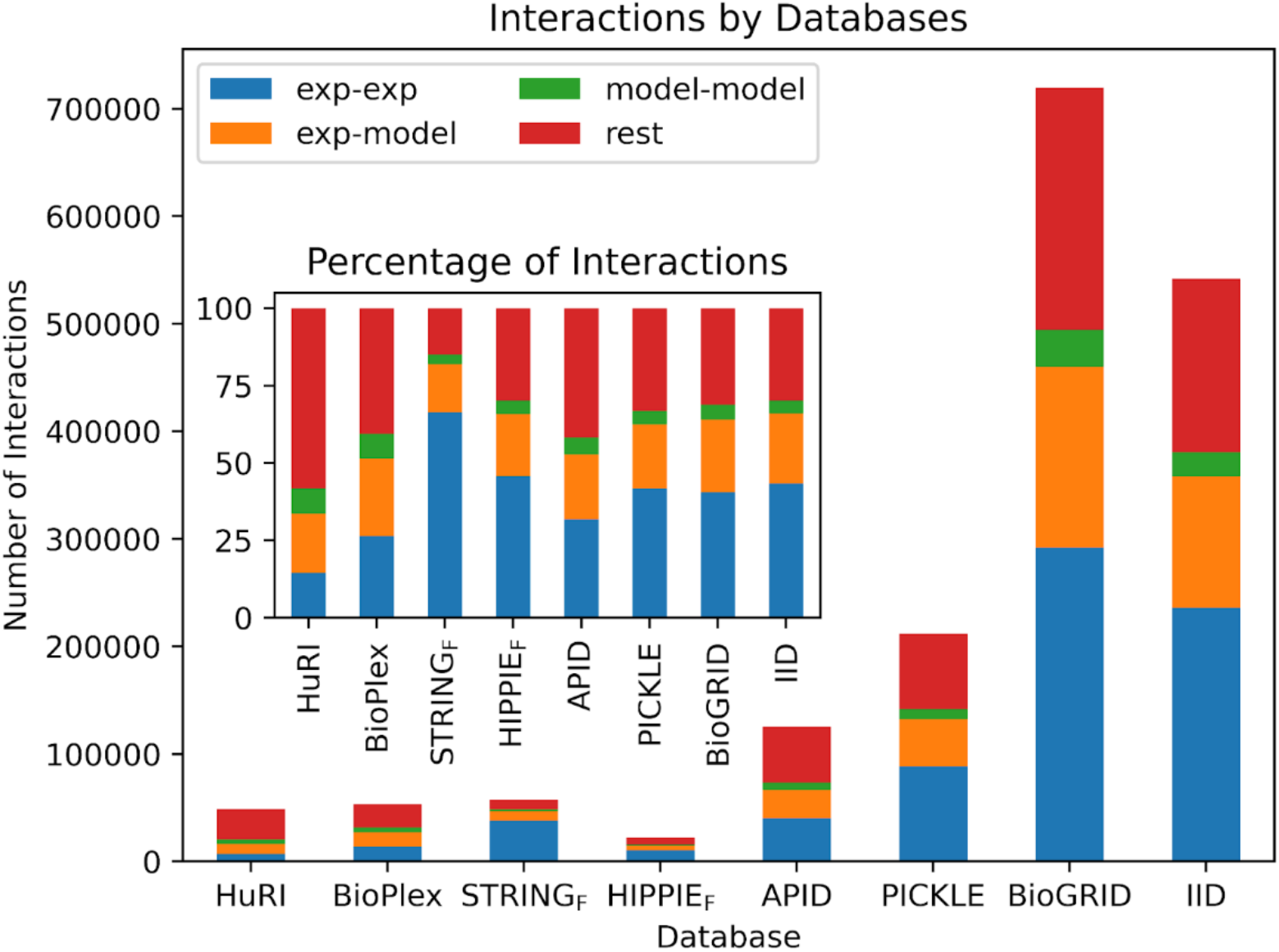
Structural coverage of protein-protein interactions in interactome databases. Blue indicates interactions where both interacting partners have *experimental* structures from PDB (exp-exp). Orange represents interactions where only one interacting partner has an *experimental* structure from PDB, and the other partner has a *model* from either ModBase, SWISS-MODEL, or AlphaFold (exp-model). Green indicates that both interacting partners do not have experimental structures but have a *model* (model-model). Lastly, red for the remaining interactions which have no structural data available (rest). The inset plot shows the same concept in terms of percentages. AlphaFold models that have 85% of their residues predicted with ≥ 70% pLDDT score are considered.

Accurate 3D modeling of the interactions in HuRI, STRING_F,_ and HIPPIE_F_ interactomes requires structurally complete monomer protein structures. In other words, given that two proteins interact, structural knowledge for this protein-protein interaction can be obtained on the availability of the structures of both proteins. To assess the completeness of each interacting pair, proteins in HuRI, STRING_F_ and, HIPPIE_F_ interactomes were labeled as high coverage (HC) or low coverage (LC) along with the name of the data sources: PDB, SWISS-MODEL, or ModBase. On the other hand, AlphaFold usually predicts the whole protein structure, yet the prediction quality of each structure, even each residue in a structure, is different. Therefore, rather than using HC and LC metrics, we preferred high accuracy (HA) and low accuracy (LA) metrics for the AlphaFold structures. **Figure 6** shows a snapshot of the available structures in different databases. As stated in previous sections, we denoted 50% or higher coverage as HC; the rest as LC; 85% of the structure covered with 70% or higher accuracy for AlphaFold, is designated as HA, with the remaining predictions falling into the LA group. Numerous combinations of databases and coverages are available for an interaction because the 3D structures of each protein can be found across multiple databases with various coverages or accuracies. This is why the sum of the numbers in **Figure 6** is greater than the total number of interactions in the reference interactome. AlphaFold’s low accuracy 3D structures dominate the HuRI, STRING_F_, and HIPPIE_F_ interactomes. PDB and homology modeling databases each add roughly equal numbers of structures to modeling a small number of interactions in HuRI and HIPPIE_F_. On the other hand, the PDB and homology modeling databases contain many 3D structures with high coverage that can be used to model the interactions in STRING. Only 12.97% of the HuRI and 32.94% of the HIPPIE_F_ interactomes contained protein partners with high accuracy or high coverage structures, while this percentage increased to 73.62% for the STRING_F_ interactome. Out of 117,897 interactions in total, 47,431 (40.23%) interactions have both protein partners with high accuracy or high coverage structures.

**Figure 6.**
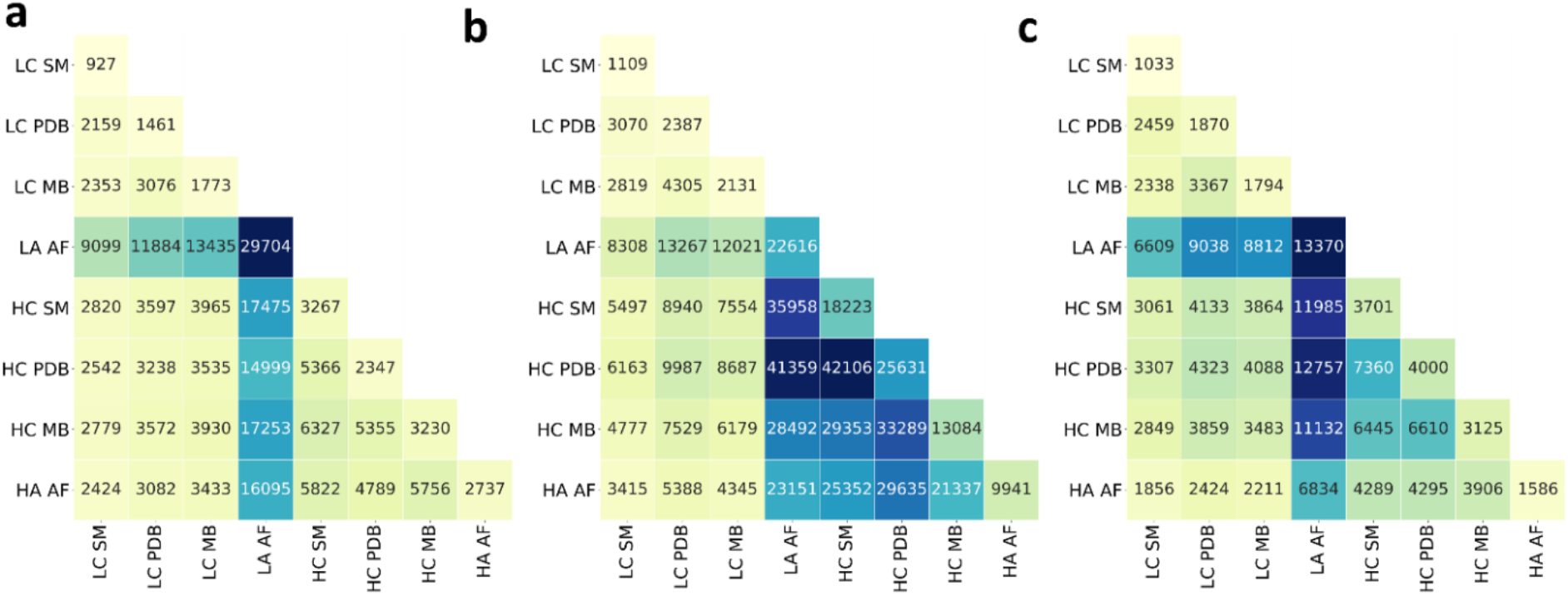
3D Structure modeling assessment of “Protein 1-Protein 2” pairs in reference interactomes in. (a) HuRI (b) STRING_F_ (c) HIPPIE_F_. x-y axes labels show the 3D coverage or accuracy label of the databases. LC: low coverage, HC: high coverage, HA: high accuracy, LA: low accuracy, MB: ModBase, SM: SWISS-MODEL, AF: AlphaFold. The number in each cell indicates how many “Protein 1-Protein 2” pairs of the reference interactome can be potentially constructed by using mentioned sources with given labeled coverage or accuracy.

Some of the proteins are highly studied; these usually correspond to disease-related proteins. Investigation of important proteins such as disease-related, COSMIC CGC [48], and drug-targets in interactome databases shows that except for HuRI, interactomes contain ≥ 50% of COSMIC CGC and disease-related genes (**Figure 7**). Even though the number of interactions in the HIPPIE_F_ dataset is less than HuRI, it is enriched in important genes. The coverage of drug targets, however, is often less extensive across most interactomes. When our selected reference interactomes are combined, drug target coverage increases by up to 69% while others surpass 75%.

**Figure 7.**
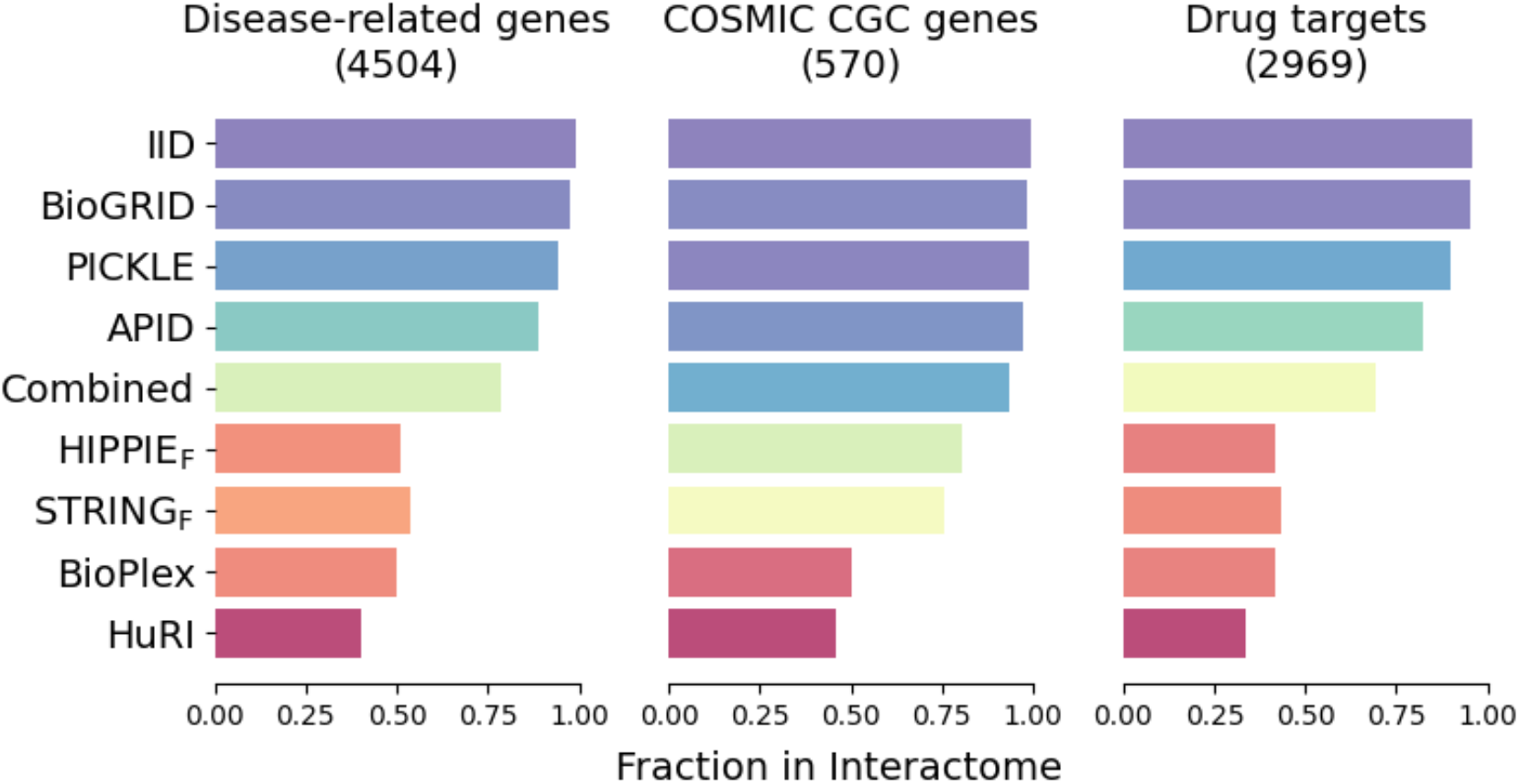
Investigation of important genes in interactome databases. Fraction of disease-related, COSMIC CGC, and drug targeting genes are investigated for all interactome databases. The term “combined” represents merged interactomes of HIPPIE_F_, STRING_F,_ and HuRI.

## Discussion

We comprehensively analyzed the current structural knowledge of the human proteome and interactomes (HuRI, STRING_F_, and HIPPIE_F_) by integrating experimental protein structures from PDB and predicted structures from ModBase, SWISS-MODEL, and AlphaFold. Our results showed that combining several resources improved the structural coverage of the human proteome and interactomes. Predicted protein structures bring an orthogonal layer of information toward having a more comprehensive structural proteome with varying degrees of accuracy. The availability of an experimental model significantly impacts the computational accuracy of the structure prediction. Given that experimental structure determination is more difficult for the antibody class of proteins due to their long loop regions [49], our data demonstrate that the prediction ability is indeed restricted for these proteins. Proteome-level analysis shows we are still halfway to a structurally complete human proteome.

Integrating the structural data mapped to reference interactome databases at the interactome level reveals that STRING_F_ has the highest PDB coverage while HuRI has the lowest. This suggests that STRING may be a valuable resource for researchers seeking information about interactions involving experimental structures. While the number of proteins with PDB structures is similar between interactome databases, the number of interactions differs significantly, which might be attributed to the limited sensitivity of Y2H. Therefore, additional computational prediction methods, such as template-based or template-free docking and deep-learning-based methods, complement experimental methods as they capture interactions that Y2H or other experimental techniques may miss.

We also highlight that homology models show better interaction coverage than AlphaFold’s accurately predicted models. This implies that homology models may continue to be a useful tool for predicting PPIs when experimental data is lacking and may complement the capabilities of AlphaFold. Furthermore, we emphasize that there is minimal overlap between interactome databases. Consequently, depending solely on a single database may result in incomplete or biased results, as each database may have its strengths and limitations regarding coverage. Thus, integrating data from multiple databases can provide a more comprehensive and reliable picture of PPIs.

It has also been shown that protein pairs with high accuracy or high coverage structures constitute a small portion of interactions in HuRI, a larger portion in HIPPIE_F_, and the majority of interactions in STRING_F_. HuRI is reported as a less biased interactome containing genes that belonged to uncharted regions previously [15]. Such a characteristic might be the reason behind its lower coverage of interacting proteins than other interactomes. Considering this, bias toward well-studied proteins may impact the structural coverage of the interactome because less-studied proteins may have fewer interactions reported in the interactome. Despite the increased coverage provided by computational methods, the number of high accuracy/coverage interactions remains low compared to all existing interactions. This highlights the need for continued efforts in both experimental and computational methods to improve our understanding of protein-protein interactions and the structural coverage of the human interactome.

## Methods

### Reference Proteome Analysis

A recent version of the *UP000005640* was obtained from the UniProtKB proteome database (UniProt release 2022_04). This text-based data contains information for 20,360 reviewed human proteins. First, protein names, including keywords such as “putative” or “uncharacterized” are eliminated from the initial proteome dataset. Second, only the proteins with proven existence belonging to experimental evidence at the protein level and experimental evidence at transcript level classes are kept in the dataset. Last, proteins with less than 30 amino acids are removed from the dataset. After the three-step filtering, the final reference proteome dataset contained 18,401 proteins. Domain information was incorporated into this dataset from the PFAM database cross-referenced in UniProt. We considered a protein “multi-domain” if multiple PFAM IDs are assigned to its UniProt ID.

#### Protein Data Bank (PDB) coverage

We retrieved all PDB structures cross-referenced in UniProt to obtain the structural proteome and its coverage (as of 2022-06-01). Downloadable UniProt data only contains the identifiers of the PDB but not respective chain identifiers per entry. Some PDB coordinate files might not represent the entire structure but partial structure. Besides, due to experimental limitations, residues might have missing coordinates within the covered part. These conditions have to be considered to make an exact coverage calculation. We parsed all mm/ CIF files by using Biopython’s Bio.PDB package [50] to find the protein chains that match UniProt identifiers in the reference proteome. In total, we parsed 128,696 protein chains for 7,376 unique human proteins. Calculations run on Koc University High-Performance Computing (KUACC HPC) cluster. In the end, we have an exact begin-end residue range for each PDB chain corresponding to the original UniProt sequence and missing residues that fall into that region. The corresponding PDB file is not considered for further analysis if the resulting region does not contain at least 30 consecutive amino acids. This coordinate to UniProt sequence mapping allowed us to see the exact sequence coverage of each entry in our reference proteome. To eliminate redundancies, we removed duplications representing identical residues and excluded missing coordinates only once if there were any repeats in another coordinate file. For all PDB coverage calculations, Equation 1 is used.

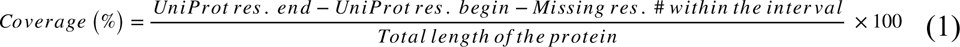

#### AlphaFold (AF) coverage

The DeepMind team released structures predicted by AlphaFold v.02 [51] in Oct 2022. Predictions for UP000005640 proteome downloaded from AlphaFold Protein Structure Database (https://ftp.ebi.ac.uk/pub/databases/alphafold), which Deep Mind and EMBL-EBI developed. There were 23,391 predictions deposited for the reference proteome. After eliminating the predictions previously labeled as unreviewed by UniProt, we ended up with 20,315 reviewed/canonical protein predictions. AlphaFold predicts the proteins as fractions whose amino acid count exceeds 2,700. Fractions are deposited as separate coordinate files and mostly overlap regarding residue identifiers. Those predictions for 207 unique protein entries are also eliminated. All analyses were applied to the remaining 20,038 protein predictions. Confidence in the AlphaFold predictions is measured by pLDDT, a per residue accuracy estimate on a scale from 1 to 100. A pLDDT score ≥ 70% is declared as an indicator of a good backbone prediction [40]. Therefore in our analysis, we applied a structural constraint. We only assume “accurately predicted” if a protein’s 85% of the residues predicted with ≥ 70% pLDDT score.

#### Homology modeling coverage

We have utilized two widely known homology modeling databases: SWISS-MODEL and ModBase. We downloaded homology models for reference proteome based on the UniProtKB release 2022_04 from the SWISS-MODEL Repository. We eliminated models that are not present in the filtered human reference proteome and have less than 30 amino acids. Then, we filtered the models based on their QMeanDisCo Global score, a composite scoring function that estimates model quality [52]. We selected models with a QMeanDisCo Global score greater than 0.7 as they are considered confident models by the providers. For ModBase, we downloaded the file named *modbase_models_all-latest.xz* with a last modified date of 28/02/2014 from the downloads section. Like before, we eliminated models that are not present in the filtered human reference proteome and have less than 30 amino acids. We selected models with a ModPipe Quality Score (MPQS) ≥ 1.1 if available; or if MPQS is not available, sequence identity ≥ 30% and a model score (GA341) ≥ 0.7 were selected as good quality models MPQS is a composite quality score that includes e-value, z-Dope, GA341, coverage, and sequence identity to the template [53]. However, the total length of some proteins given in ModBase was greater than the length reported by UniProt. This caused the coverage for those proteins to be more than 100%. Hence, we decided not to use the models of these proteins. Also, we selected the model with a higher quality score for both databases if there were models with precisely the same amino acid start and end positions. We have reported residue-based and protein-based coverage results for these models. For residue-based coverage calculations, a simplified version of Equation 1 is used where the missing residues are not considered because of the nature of homology models.

To understand the contribution of homology modeling to experimentally-determined protein structures, we further eliminated the models that cover the identical residues with available PDB structures. We have calculated the coverage of the proteome again by only considering the residues modeled by homology databases and not by PDB. This way, we could observe the actual contribution of homology models regarding structural coverage. We have utilized the former case where we do not eliminate models concerning PDB data for the analysis of interactome coverage.

#### Structural coverage of the protein classes

In this study, we used functional classification data of the proteins from the Human Protein Atlas [54]. Five main protein classes: membrane proteins, transcription factors, antibodies, and transporter proteins, are investigated in terms of their structural coverage either experimentally (PDB) or computationally (AlphaFold). Only the proteins in our reference proteome are filtered from the Human Protein Atlas data. As AlphaFold produces whole protein predictions, its coverage is evaluated in terms of prediction accuracy rather than the count of the protein residues predicted. In this manner, average counts of the proteins in any class are calculated by dividing the number of proteins with a 3D structure by the protein count within the class.

### Analysis of interactome databases

We reviewed interactome databases currently available online, including HuRI [15], STRING [9], BioPlex [5], BioGRID [14], HIPPIE [10], Interactome INSIDER [3], Interactome3D [4] hu.MAP [16], IID [7], APID [6] and PICKLE [12]. An overview of these databases can be found in Supplementary Table 1. We downloaded the protein-protein interactions from HuRI, STRING, BioPlex, BioGRID, HIPPIE, IID, APID, and PICKLE databases to investigate the number of PPIs available (as of December 5, 2022). We downloaded the HuRI.tsv file of the 2021 publication from http://www.interactome-atlas.org/download. STRING physical interaction dataset named “9606.protein.physical.links. detailed.v11.5.txt” was downloaded from https://stringdb.org/cgi/download?sessionId=beWWyYG7kyNt&species_text=Homo+ sapiens and rescoring have been performed using the Python script from https://stringdb-static.org/download/combine_subscores.py, which STRING creators provide. We adjusted the scoring to only utilize the scores of experimental and database channels and calculated a new combined score. We selected models with a combined score of experimental and database channels greater than 0.7. For BioPlex, we downloaded the latest dataset BioPlex 3.0 Interactions (293T Cells) from https://bioplex.hms.harvard.edu/interactions.php. For HIPPIE, we downloaded the current release (v2.3) from http://cbdm-01.zdv.uni-mainz.de/

<u>∼mschaefer/hippie/download.php</u>. We filtered this dataset which collects PPIs from studies with a wide range of experimental methods so that we have only the binary PPI detection methods which are two-hybrid, atomic force microscopy, and fluorescent resonance energy transfer [1]. Then, we set the quality threshold at 0.73 (high quality) and removed interactions smaller than this cutoff value. The latest version (March 2021) of APID data was downloaded from http://cicblade.dep.usal.es:8080/APID/init.action where level 2 was selected, and inter-species interactions were filtered out. In the case of PICKLE, we downloaded the current release (v3.3) normalized at the protein (UniProt) level and cross-checked from http://www.pickle.gr/Downloads#HUMAN-3-3. BioGRID release 4.4.216 with a last modified date of November 29th, 2022, was downloaded from https://downloads.thebiogrid.org/ <u>File/BioGRID/ReleaseArchive/BIOGRID4.4.216/BIOGRID-ORGANISM-4.4.216.tab3.zip</u> where *Homo sapiens* interactions were extracted later, and physical experimental system type was selected. Lastly, we downloaded the IID dataset (v2021_05) named human_annotated_ PPIs.txt.gz from the downloads section of http://iid.ophid.utoronto.ca and filtered the dataset by selecting interactions with evidence types that included experiments. We converted any different protein identifiers to UniProt identifiers for standardization and removed redundant protein-protein interactions. We filtered the datasets for the proteins present in our reference proteome. Then, we found the number of unique proteins and unique interactions in each interactome database, the number of unique proteins with at least one available structure in PDB, and the number of unique proteins with at least one model in SWISS-MODEL, ModBase, and AlphaFold databases. Furthermore, we found a number of unique interactions in which both interacting pairs had a structure and/or a model.

#### Gene Ontology (GO): Cellular component analysis of interactome databases

For this part, we focused only on the proteins available in STRING to reveal the reason behind the difference in the interaction counts between interactome databases. We collected the PFAM IDs of these proteins and found UniProt entries in the reference proteome containing at least one PFAM domain. . A common pattern in terms of frequent GO cellular component annotations [55], ribosome (GO:0005739), and/or mitochondrion (GO:0005840) are detected for this protein subset. Then, we checked the existence of such proteins with these annotations in STRING_F_, HuRI, and HIPPIE_F._

#### Investigation of important genes in interactome databases

We investigated the coverage of disease-related, COSMIC CGC, and drug-targeting genes in interactome databases. For disease-related genes, we downloaded variant summary data from Clinvar [56] (last modified date: 2023-01-21) and filtered them to take genes in assembly GRCh38 and have ‘Pathogenic’ clinical significance. For COSMIC CGC genes, we downloaded cancer gene census data from COSMIC (version 97) and restricted them to Tier 1 genes [48]. Lastly, for drug targets, we utilized the complete target data from IUPHAR/BPS Guide to Pharmacology (2022.4 version) [57]. We filtered all datasets for the proteins present in our reference proteome. Then, we calculated the fraction of these important genes in interactome databases to find their coverage.

## Notes

### Competing Interest Statement

The authors have declared no competing interest.

