## Supplementary Figure 1 for "Structural Coverage of the Human Interactome"

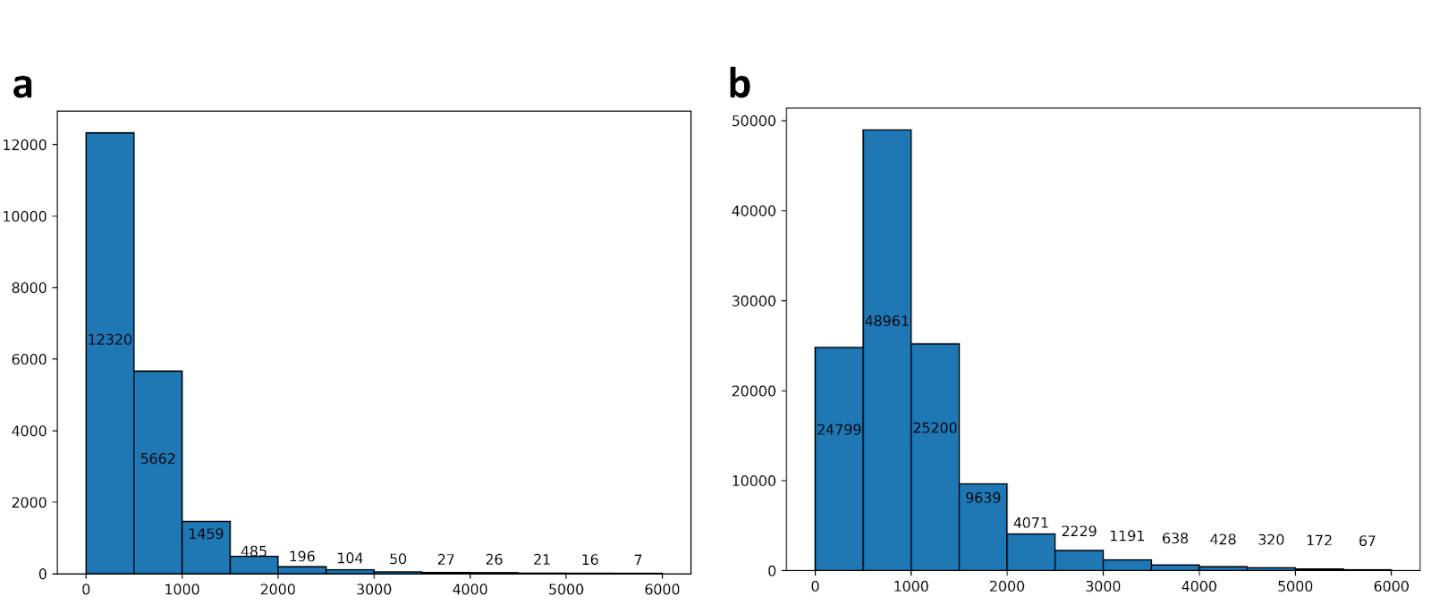
 **Supplementary Figure 1.** In histograms the y-axis shows the protein counts and x-axis shows protein lengths. **(**a) Protein sequence length distribution of the human reference proteome (b) Interaction sequence length distribution of the reference interactomes.

**Supplementary Table 1.** Overview of interactome databases.

| **Database** | **# of organisms** | **Data source** | **Stats (*homo sapiens*)** | **Pros** | **Cons** | **Website** |
| --- | --- | --- | --- | --- | --- | --- |
| **HuRI** | 1 | Yeast Two-Hybrid (Y2H) | 52,569 PPIs | Extensive search space (~90% of the protein-coding genome), gives binary PPIs | Many PPIs may not be detected because the sensitivity of the Y2H method is limited. | <http://www.interactome-atlas.org/> |
| **Interactome3D v2020_05** | 18 | IntAct, DIP, InnateDb, BioGRID, BINDTranslation, HPRD, 3did | 136,329 PPIs [structural data for 8,359 interactions, models for 7,625 interactions, 120,345 structureless] | Incorporates structural information into PPI networks, provides structurally annotated pathways and 3D coordinates | Does not predict whether two proteins interact | <https://interactome3d.irbbarcelona.org/> |
| **PrePPI v1.2.0** | 2 | MIPS, DIP, IntAct, MINT, HPRD, BioGRID and prediction | 1.35 million PPIs [372,545 high confidence PPIs (> 0.5)] | integrates predicted & experimental PPIs; combines structural and non-structural info to predict PPIs | high number of false positives | <https://honiglab.c2b2.columbia.edu/PrePPI/> |
| **Interactome INSIDER** | 8 | DIP, IntAct, MINT, BioGRID, iRefWeb, HPRD, MIPS | 121,575 PPIs | map of interaction interfaces, disease mutation exploration, prediction of interfaces | disordered interfaces are not included in training set | <http://interactomeinsider.yulab.org/> |
| **BioPlex v3.0** | 1 | AP-MS in 293T and HCT116 cells | 118,162 PPIs (293T); 70,966 PPIs (HCT116) | the most comprehensive experimentally derived model, two context-specific interaction networks | AP-MS cannot detect some PPIs and resultant PPIs are not always binary PPIs | <https://bioplex.hms.harvard.edu/> |
| **hu.MAP v2.0** | 1 | Integration of over 15,000 mass spectrometry experiments using a machine learning framework | 57,148 PPIs | uncharacterized genes are better covered, reduced false positive and negatives | MS methods are unable to detect some PPIs, the website is less developed | <http://humap2.proteincomplexes.org/> |
| **IID v2021_05** | 18 | BCI, BIND, BioGRID, DIP, HPRD, I2D, InnateDB, IntAct, MatrixDB, MINT, ortolog ve tahmin | 1,209,534 PPIs [560,628 experimental, 92,600 orthologous, 660,503 predicted] | context-specific; annotations such as tissue, disease, cellular localization, duration, druggability, directionality, membership in complexes, conservation across species, curation opted towards reducing false negatives | the search part of the website is less practical | <http://iid.ophid.utoronto.ca/> |
| **STRING v11.5** | 14,094 | BIND, DIP, GRID, HPRD, IntAct, MINT, PID for exp data; Biocarta, BioCyc, GO, KEGG, Reactome for curated data & prediction | 11,938,498 PPIs (functional included), 1,991,832 physical PPIs | evidence from 7 different sources and scores for each, physical interactions can be downloaded separately, website interface is convenient, offers multiple download options |  | <https://string-db.org/> |
| **BioGRID v4.4.216** | 81 | experimental [36,460 pubs for human] | 1,188,899 PPIs | gives experimental PPIs, protein-molecule interactions and PTMs, has themed curation projects | does not guarantee that the interaction is direct and physical | <https://thebiogrid.org/> |
| **HIPPIE v2.3** | 1 | IntAct, MINT, BioGRID, HPRD, DIP, BIND, MIPS, Bell09, 11 studies | ~831,933 PPIs | confidence score is optimized and performs well; offers functional, tissue, impact, and disease-specific contexts | When the network is visualized on the website, the interaction between non-input protein proteins cannot be analysed | <http://cbdm-01.zdv.uni-mainz.de/~mschaefer/hippie/> |
| **APID v2021_03** | >1,100 | BioGRID, DIP, HPRD, IntAct, BioPlex | 667,805 PPIs  [154,955 binary PPIs] | possibility to download binary PPIs at different levels of experimental evidence, curation is said to be reducing false positives | visualizing a network of more than 200 proteins on the website is CPU-intensive | <http://cicblade.dep.usal.es:8080/APID>/ |
| **PICKLE v3.3** | 2 | BioGRID, HPRD, IntAct | 218,025 PPIs | covers ~80% of the proteome, tries to find interactions with high probability of being direct by three-step filtering | no options to download network visualization | <http://www.pickle.gr/> |
